# Mapping the epitopes of the Human Leukocyte Antigen E antibodies 3D12 and 4D12

**DOI:** 10.1101/2022.05.27.493692

**Authors:** Simon Brackenridge, Persephone Borrow, Andrew J. McMichael

**Affiliations:** Centre for Immuno-Oncology, Nuffield Department of Medicine, Old Road Campus Research Building, University of Oxford, Oxford, OX3 7DQ, United Kingdom

## Abstract

The commonly used commercially available antibodies 3D12 and 4D12 recognise the human leukocyte antigen E (HLA-E) protein. 3D12 is known to exhibit minimal cross-reactivity with classical HLA-Ia allotypes and we confirm that this is also the case for 4D12. These antibodies bind different epitopes on HLA-E, and differ in their ability to recognise alleles of the major histocompatibility complex E (MHC-E) proteins of rhesus and cynomolgus macaques. Using hybrids of different MHC-E alleles, we have mapped the regions that are critical for the binding of these two antibodies. 3D12 recognises a region on the alpha 3 domain that is unique to HLA-E, on the opposite side to that bound by β2-microglobulin, while 4D12 recognises the start of the alpha 2 domain, adjacent to the C terminus of the presented peptide. Knowledge of the binding sites of these two antibodies will facilitate selection of the best antibody for a given application, and inform interpretation of the resulting data.

## INTRODUCTION

In contrast to the highly polymorphic classical major histocompatibility complex class Ia (MHC Ia) molecules, members of the class Ib family (MHC-E, -F, and -G) exhibit significantly less polymorphism^1^. For the human MHC-E molecule, also known as human leukocyte antigen-E (HLA-E), there are 302 functional alleles, encoding 122 distinct proteins (International Immunogenetics HLA Database, version 3.48.0). Despite this apparent diversity, only two forms of HLA-E predominate (HLA-E*01:01 and HLA-E*01:03), and the 82 and 84 alleles (respectively) encoding them appear to be under balancing selection^2^. As a result, diversity in HLA-E is effectively restricted to a single polymorphism at position 107 (arginine in HLA-E*01:01 and glycine in HLA-E*01:03). This polymorphism lies outside of the peptide binding groove and affects stability of the HLA-E/β2-microglobulin/peptide complex, resulting in higher cell surface expression of HLA-E*01:03^3^.

HLA-E preferentially presents conserved nonamer peptides derived from the signal sequences of MHC Ia proteins and HLA-G^4–6^. HLA-E in complex with this peptide (which typically has the sequence VMAPRT[V/L][V/I/L/F]L, VL9) is recognised by CD94/NKG2 receptors on natural killer (NK) cells and a subset of CD8^+^ T cells^7–9^. In addition to presenting this self-peptide, there has been a growing realisation that HLA-E also presents peptides from bacterial and viral pathogens^10^. Such HLA-E-restricted T cell responses are rare compared with those recognising peptides presented by MHC Ia, but exceptions are known. One third of the CD8^+^ T cells induced by a rhesus CMV-vectored SIV vaccine that enables ∼55% of vaccinated rhesus macaques to clear infection following challenge with SVImac239^11, 12^ are restricted Mamu-E, the rhesus orthologue of HLA-E^13^. The protection conferred by this vaccine appears to depends on these Mamu-E-restricted responses, and not the MHC class II-restricted CD8^+^ T cells that comprise the remainder of the broad and atypical vaccine response^14^. This has raised the prospect of a HIV-1 vaccine that induces protective HLA-E-restricted CD8^+^ T cells, and there is great interest in exploiting HLA-E for vaccine and immunotherapy strategies for infectious diseases and tumours^15–19^.

With increased focus on HLA-E, it is important that the available HLA-E antibodies are fully characterised. We^20^, and others^21^, have shown that some commercially available HLA-E antibodies are not truly specific. However, the most commonly used HLA-E antibody, 3D12^9^, has minimal cross-reactivity with MHC Ia^20^. The laboratory that isolated 3D12 later isolated a second HLA-E antibody, 4D12^22^. Although 4D12 was reported not to bind a panel of 60 representative cell lines, we are not aware of any comprehensive analysis of its cross-reactivity with MHC Ia. In terms of MHC-E reactivity, however, 4D12 differs from 3D12 by staining cells expressing Mamu-E*02:04^13^ and the cynomolgus macaque orthologue Mafa-E*02:01:02^23^.

In addition to understanding the repertoire of proteins that these two antibodies can bind, knowledge of the locations of their epitopes will allow determination of whether they are suitable for use in blocking engagement with particular receptors, or whether biotinylation of exposed lysine residues in or near their binding sites could prevent antibody binding. Knowledge of the epitopes may also be critical to understand what external factors could alter interaction of the antibodies. It has recently been shown that the binding of the pan-MHC class I antibody W6/32 can, in certain situations, be influenced by the lipid composition of the plasma membrane^24^. Membrane compositions can differ significantly between cell types and activation states^25^, so knowing where an antibody binds on a cell surface protein may be essential to the correct interpretation of staining patterns.

3D12 and 4D12 recognise different regions of HLA-E^22^, but these have never been properly mapped. One attempt to map the 3D12 epitope suggested two regions on the HLA-E extra-cellular domain (ECD)^26^ that are conserved in Mamu-E*02:04. As this allotype is not recognised by 3D12, these sequences alone cannot comprise the epitope. A later attempt to map the 3D12 epitope using overlapping peptides spanning the HLA-E protein was also unsuccessful, and it was concluded that the 3D12 epitope has a moderate conformationally-dependent component^27^.

To map the epitope of 3D12, we exploited the fact that this antibody binds HLA-E*01:03 but not Mamu-E*02:04 and. Staining of cells expressing hybrids of these two proteins revealed a region of sequence unique to HLA-E on the alpha 3 domain that was critical for 3D12 binding. For 4D12, staining of cells expressing hybrids of Mamu-E*02:04 and Mamu-E*02;16 (which is not recognised by 4D12) allowed us to map the 4D12 epitope to the N-terminal end of the MHC-E alpha 2 helix, adjacent to the C terminus of the presented peptide. Finally, we confirmed that 4D12, like 3D12, does not cross-react with MHC Ia.

## METHODS

### Plasmids & cloning

The single chain trimers (SCT) of Mamu-E*02:04 and HLA-E*01:03 with the VL9 peptide (VMAPRTLLL) were as previously described^13, 28^. Additional SCT of HLA-E*01:01 and Mamu-E*02:16 were created using an identical cloning strategy. Hybrids of the SCT constructs were made using restriction enzymes (PpuMI, Bsu36I, and SbfI). All plasmids were verified by sanger sequencing using an ABI 3730.

### Cell culture, transfection, staining & flow cytometry

HEK 293T cells were maintained between 10% and 90% confluency at 37°C/5% CO2 in DMEM (Life Technologies) supplemented with 10% foetal bovine serum (Sigma), and penicillin/streptomycin (50 units/ml and 50 µg/ml, respectively; Life Technologies). Transfections were carried out in 6 well plates using GeneJuice (Merck) with 1 μg of plasmid DNA per well, as per the manufacturer’s instructions. Cells were stained 24 hours post transfection as previously described^13^ with 3D12 (BioLegend), 4D12 (MBL), or 2M2 (BioLegend). Stained cells were acquired using a CyAn ADP Analyser (Beckman Coulter), and analysed using the gating strategy shown in Supplementary Figure 1A using FlowJo 10 (BD).

### Inactivation of β2-microglobulin

A β2-microglobulin-specific gRNA (5’-CGCGAGCACAGCTAAGGCCA-3’^30^, which targets the reverse strand of the first exon) was inserted into pspgRNA^31^ (a gift from Charles Gersbach, Addgene plasmid # 47108). 293T cells were co-transfected with equal amounts of this plasmid and pCas9_GFP^32^ (a gift from Kiran Musunuru, Addgene plasmid # 44719). GFP-positive cells were sorted after 48 hours on a MoFlo cell sorter (Beckman Coulter), expanded after diluting to 0.7 cells per well in 96 well plates, and 8 clones that were negative by surface staining with W6/32 (BioLegend) grown out. One clone which grew and transfected well was selected for use. Transfection of these cells with a plasmid expressing β2-microglobulin was sufficient to restore surface W6/32 staining, confirming that the loss of W6/32 staining resulted disruption of the β2-microglobulin gene (Supplementary Figure 1B).

### Characterisation of 4D12 and 3D12cross-reactivity

Cross-reactivity of 3D12 and 4D12 was assessed using LABScreen Single Antigen Class I Combi beads (One Lambda) as previously described^20^. Raw signals were corrected by subtracting the fluorescence of the negative control beads, and the binding of 3D12 and 4D12 was normalised relative to the binding of W6/32, to control for variation in the amount of correctly folded protein for each HLA allotype.

## RESULTS

### MHC-E allele reactivity of the commercially available HLA-E antibodies

To map the binding sites of the two antibodies on HLA-E we employed three alleles of MHC-E: HLA-E*01:03, Mamu-E*02:04, and Mamu-E*02:16. The sequences of the ECD of these proteins are shown in Figure 1A. As its expression is normally very low, we expressed MHC-E as single chain trimers (SCT, Figure 1B) of the MHC protein, β2-microglobulin and the VL9 peptide, linked by flexible glycine-serine linkers. Good surface expression of all three SCT was confirmed using a β2-microglobulin-specific antibody (Figure 1C, top row). 3D12 and 4D12 (Figure 1C, middle and bottom rows, respectively) both stained cells expressing the HLA-E*01:03 SCT, while 4D12 also stained cells expressing the Mamu-E*02:04 SCT. Neither antibody stained cells expressing the Mamu-E*02:16 SCT.

**FIGURE 1:**
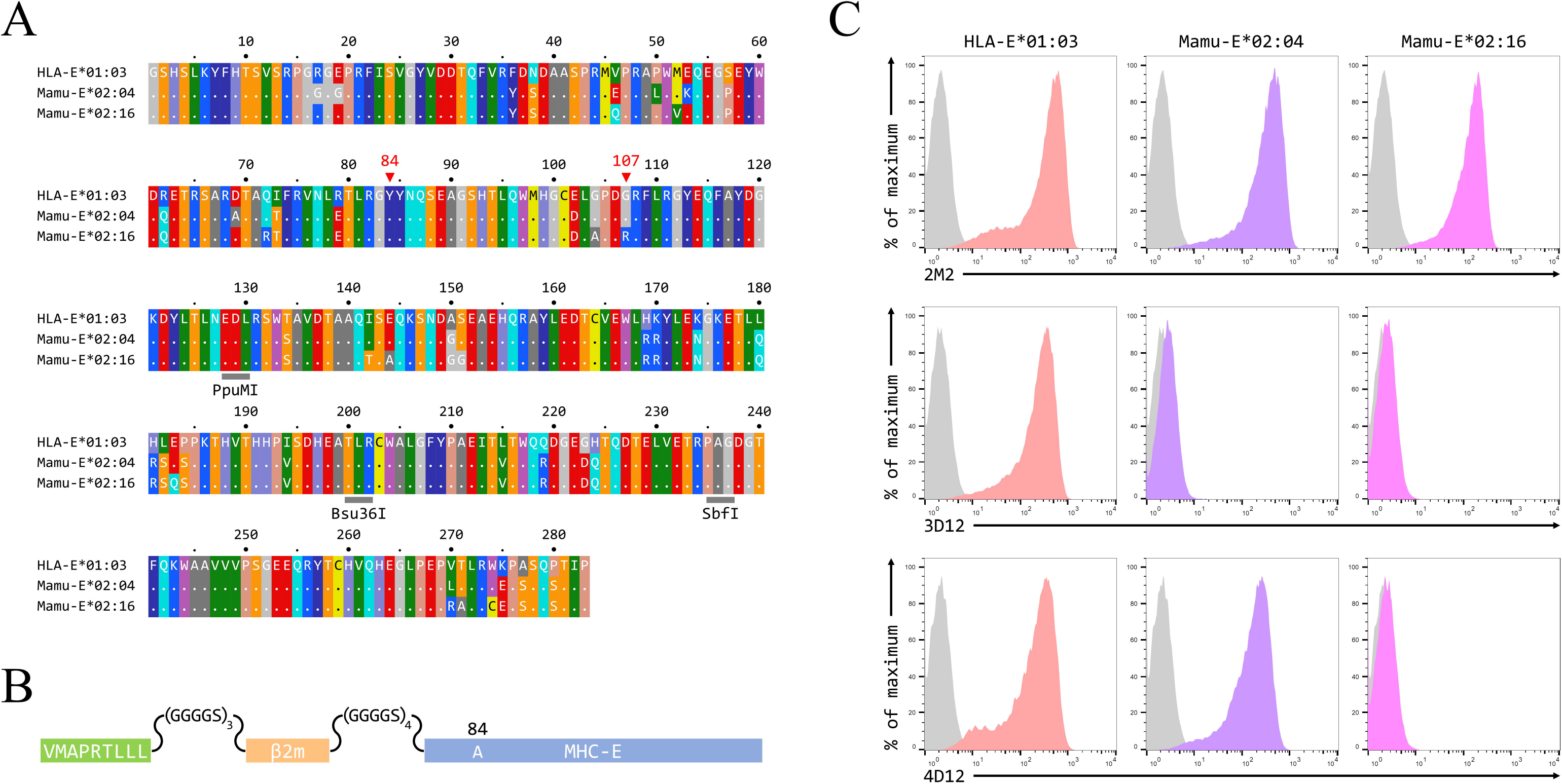
Reactivity of 3D12 and 4D12 with the alleles used in this study. **(A)** Sequences of the ECD of HLA-E*01:03, Mamu-E*02:04, and Mamu-E*02:16. Sequence identity is indicated by dots and amino acids are coloured according to the standard scheme used by RasMol. Positions 107 (the predominant polymorphism in HLA-E), and the conserved tyrosine at position 84 (mutated to alanine in the SCT to open the end of the peptide binding groove) are indicated. The approximate locations of the three restriction sites used to create the hybrids of the different alleles are also highlighted. **(B)** Schematic of the SCT constructs, which encode the peptide VMAPRTLLL, a flexible glycine-serine linker (GGGGSx3), the mature β2-microglobulin coding sequence, a second flexible linker (GGGGSx4), and the mature coding sequences of HLA-E*01:03, Mamu-E*02:04, or Mamu-E*02:16. The constructs also include the coding sequence of EGFP at the C terminus (not shown) to allow gating of transfected cells. **(C)** Representative staining with 2M2 (a β2-microglobulin-specific antibody, top row), 3D12 (middle row), or 4D12 (bottom row), of β2-microglobulin-deficient 293T cells transfected with plasmids expressing the HLA-E*01:03 SCT (first column), the Mamu-E*02:04 SCT (central column), or the Mamu-E*02:16 SCT (final column). Grey histograms show staining of mock transfected cells.

### Mapping the primary contacts of 3D12

To map the 3D12 epitope, we created a series of hybrids of the HLA-E*01:03 and Mamu-E*02:04 SCT (Figure 2A and C), using three restrictions sites shared by both alleles (PpuMI, Bsu36I, and SbfI). The ECD of these allotypes differ at 31 positions, with the hybrids having 13, 23 and 27 of these polymorphisms swapped (denoted as H13/M18 and M13/H18, H23/M8 and M23/H8, or H27/M4 and M27/H4, depending on whether the human or macaque sequence comes first). All hybrids expressed well at the cell surface (Figure 2B and D, top row), but only cells expressing the H27/M4 SCT (Figure 2B, bottom row), and M13/H18 and M23/H8 SCT (Figure 2D, bottom row), were stained by 3D12.

**FIGURE 2:**
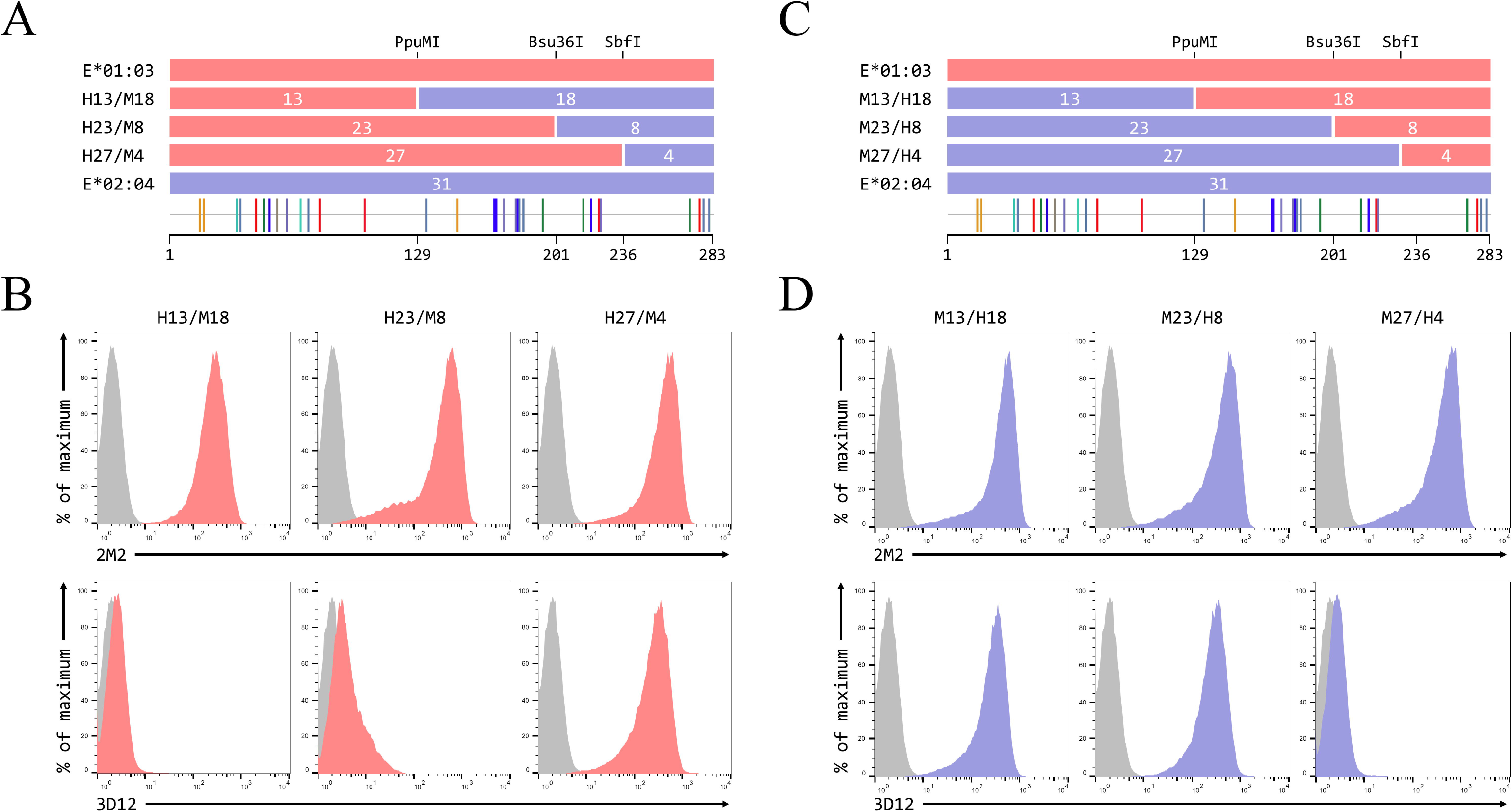
Identification of residues critical for the binding of 3D12. **(A, C)** Schematics of the ECD of the HLA-E*01:03/Mamu-E*02:04 (panel A) or Mamu-E*02:04/HLA-E*01:03 (panel C) hybrids a used to map the 3D12 epitope, which were created using the conserved PpuMI, Bsu36I or SbfI restriction sites. The highlighter plots at the bottom indicate the locations of the 31 positions that differ between the extracellular domains of these two alleles, and the number of these on either side of each restriction site (13 and 18, 23 and 8, or 27 and 4) are indicated. **(B, D)** Representative staining with 2M2 (top row), or 3D12 (bottom row), of β2-microglobulin-deficient 293T cells transfected with plasmids expressing the HLA-E*01:03/Mamu-E*02:04 (panel B) or Mamu-E*02:04/HLA-E*01:03 (panel D) hybrid SCT. Grey histograms show staining of mock transfected cells.

This suggested that the region between the Bsu36I and SbfI restrictions sites, containing the polymorphisms at positions 215, 219, 223 and 224 on the alpha 3 domain, is critical for the binding of 3D12. Across this region, Mamu-E*02:04 more closely resembles HLA-A*02:01 (and the majority of MHC Ia allotypes) than HLA-E (Figure 3A), differing only at position 215 (valine rather than leucine). In contrast, HLA-E matches MHC Ia allotypes at position 215, with the amino acids at positions 219, 223 and 224 being unique to HLA-E. To confirm this, two further hybrids were made with only these four polymorphisms swapped (H23/M4/H4 and M23/H4/M4, Figure 3B). Both SCT expressed well at the cell surface (Figure 3C, top row), but only cells expressing M23/H4/M4 stained with 3D12 (Figure 3C, bottom row). Moreover, staining of cells expressing the HLA-E*01:03 and M23/H4/M4 SCT titrated identically when decreasing amounts of 3D12 were used (Figure 3D), confirming that all sequences necessary for 3D12 to bind are present in the M23/H4/M4 hybrid.

**FIGURE 3:**
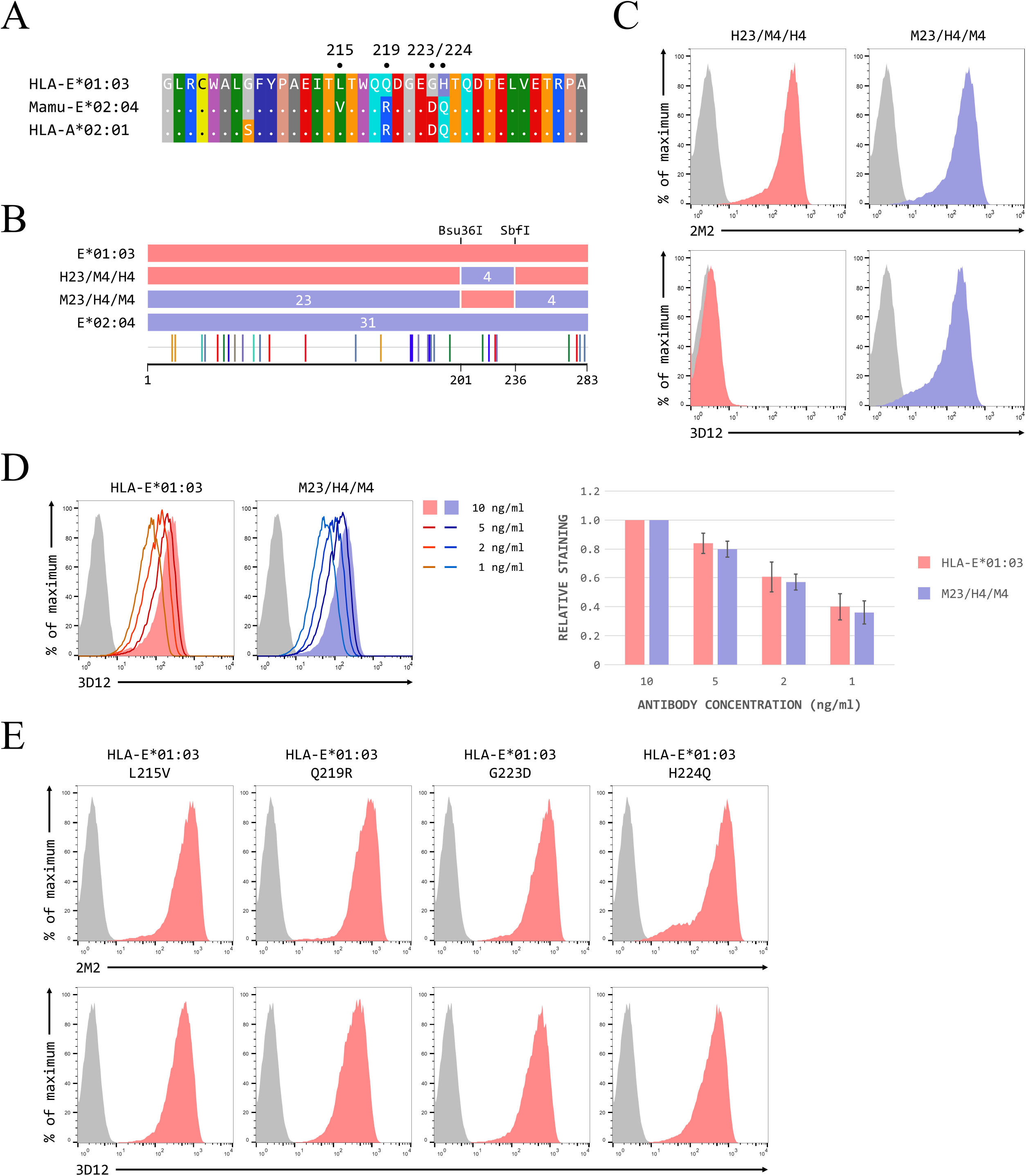
The region encompassing the polymorphisms at positions 215, 219, 223 and 224 is critical for 3D12 binding. **(A)** Comparison of the sequence between the Bsu36I and SbfI restriction sites (positions 200 to 236) of HLA-E*01:03, Mamu-E*02:04, and HLA-A*02:01. Sequence identity is indicated by dots and amino acids are coloured according to the standard scheme used by RasMol. **(B)** Schematic of the ECD of the HLA-E*01:03/Mamu-E*02:04 (H23/M4/H4) and Mamu-E*02:04/HLA-E*01:03 (M23/H4/M4) hybrids, which have the sequence between the Bsu36I and SbfI restriction sites swapped. **(C)** Representative staining with 2M2 (top row), or 3D12 (bottom row), of β2-microglobulin-deficient 293T cells transfected with plasmids expressing the H23/M4/H4 (left column) or M23/H4/M4 (right column) hybrid SCT. **(D)** Representative staining using decreasing amounts of 3D12 of β2-microglobulin-deficient 293T cells transfected with plasmids expressing the HLA-E*01:03 (left) or M23/H4/M4 (right) hybrid SCT. The graph shows the average normalised median fluorescence intensity (staining with the highest amount of antibody set to 1) for four independent repeats, and the error bars correspond to the standard error of the mean (SEM). **(E)** Representative staining with 2M2 (top row), or 3D12 (bottom row), of β2-microglobulin-deficient 293T cells transfected with plasmids expressing the HLA-E*01:03 SCT with positions 215 (first column), 219 (second column), 223 (third column), or224 (fourth column) individually mutated to the amino acid present in Mamu-E*02:04. In panels C, D and E, grey histograms show staining of mock transfected cells.

To determine their individual contributions to 3D12 binding, we mutated positions 215, 219, 223 and 224 in the HLA-E*01:03 SCT, but no significant change in 3D12 staining was observed for cells expressing any of these mutants (Figure 3E). This may mean that loss of any one contact site is not sufficient to substantially affect antibody binding, or may reflect the fact that the 3D12 epitope is at least partially dependent on the conformation of the protein, and the contribution of the amino acids at these positions may be to the overall conformation of the epitope rather than direct interaction with the antibody.

In summary, our results locate the epitope of 3D12 on the alpha 3 domain of HLA-E, in the vicinity of positions 215–224. Further structural work will be required to identify exactly which residues in this region contact the antibody, and what effects the polymorphisms present in Mamu-E*02:04 have epitope conformation and/or interaction of 3D12.

### Mapping the primary contacts of 4D12

To map the epitope of 4D12, we created hybrids of Mamu-E*02:04 and Mamu-E*02:16 using the PpuMI restriction site within codons 128–131 of the ECD. The ECD of these alleles differ at 17 positions (10 of which are upstream of the hybrid junction), and the hybrids were designated 04-16 or 16-04, depending which Mamu-E allotype comes first (Figure 4A). Both hybrid SCT expressed well at the cell surface (Figure 4B, top row), but only cells expressing the 16-04 hybrid with 4D12 (Figure 4B, bottom row). Thus, the 4D12 epitope must include one or more of the 7 polymorphisms (142, 144, 151,183, 270, 271 and 274) downstream of the hybrid junction. The region encompassing the final three polymorphisms (270, 271 and 274) was discounted as the location of the 4D12 epitope, being conserved in HLA-A*02:01, to which 4D12 does not bind (Supplementary Figure 1C). Similarly, lack of conservation between HLA-E and Mamu-E*02:04 around the position 183 polymorphism suggested this region was unlikely to be the 4D12 epitope.

**FIGURE 4:**
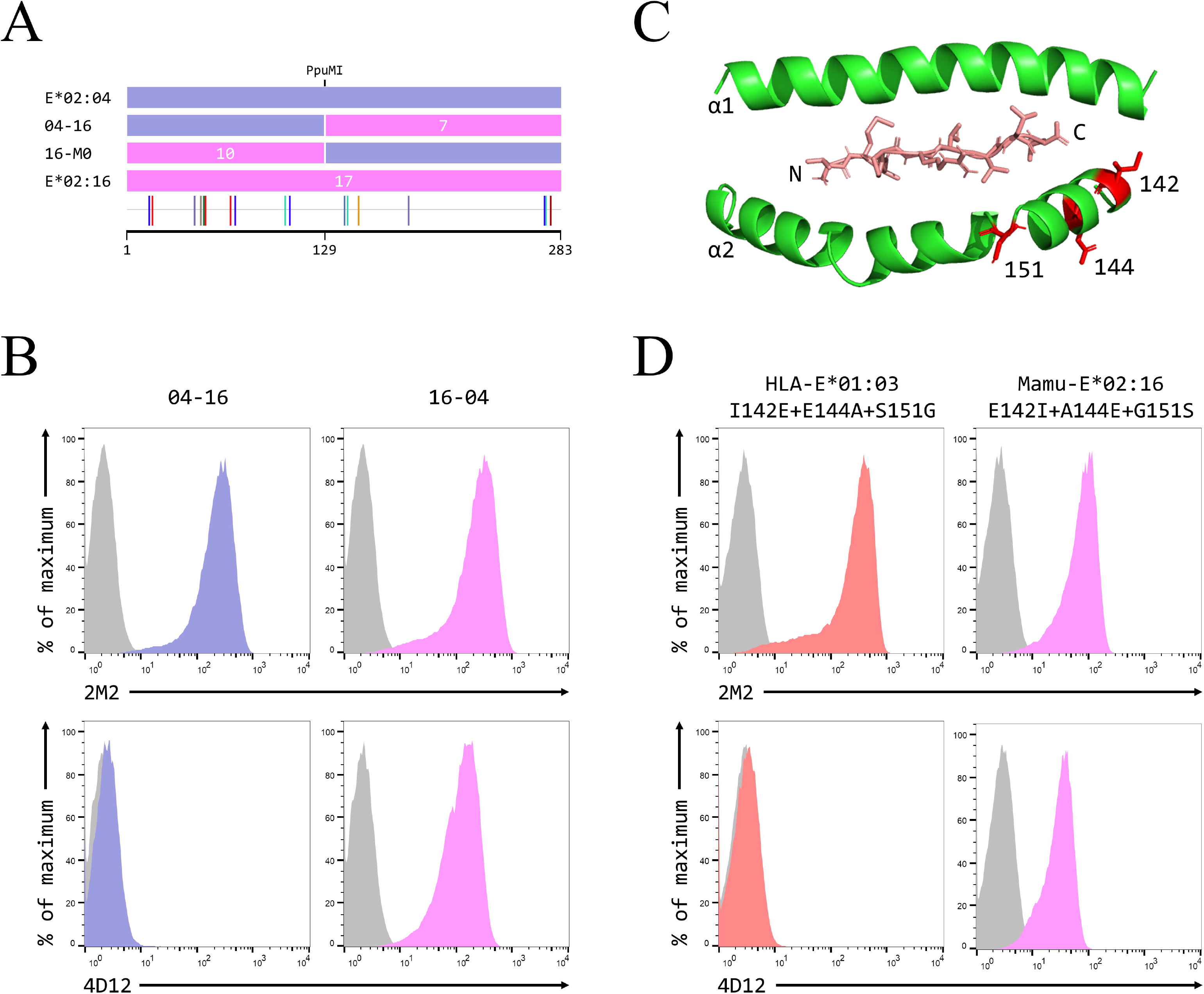
Identification of residues critical for the binding of 4D12. **(A)** Schematic of the ECD of the hybrids of Mamu-E*02:04 and Mamu-E*02:16 used to map the region bound by 4D12, which were created using the PpuMI restriction site. The highlighter plot at the bottom indicates the locations of the 17 positions that differ between the ECD of these two alleles, 10 of which lie upstream of the hybrid junction. **(B)** Representative staining with 2M2 (top row), or 4D12 (bottom row), of β2-microglobulin-deficient 293T cells transfected with plasmids expressing the 04-16 (left column) or 16-04 (right column) hybrid SCT. **(C)** Location of the three polymorphisms (positions 142, 144 and 151) most likely to encompass the location of the 4D12 epitope. The VL9 peptide is shown in stick form, while the alpha 1 and alpha 2 domains are shown as ribbons, with the side chains of the relevant polymorphic positions also included. **(D)** Representative staining with 2M2 (top row), or 4D12 (bottom row), of β2-microglobulin-deficient 293T cells transfected with plasmids expressing the HLA-E*01:03 SCT with positions 142, 144 and 151 mutated to the amino acids present in Mamu-E*02:16 (left column), the Mamu-E*02:16 SCT with positions 142, 144 and 151 mutated to the amino acids present in HLA-E*01:103 (right column). In panels B and D, grey histograms show staining of mock transfected cells.

Therefore, we focused our attention on the positions 142, 144 and 151. The first two of these lie at the N-terminal end of the alpha 2 helix, adjacent to the C terminus of the presented peptide, while 151 lies between the alpha 2-1 and 2-2 helices (Figure 4C). Combined mutation of all three positions in the HLA-E*01:03 SCT (to the residues present in Mamu-E*02:16) did not affect cell surface expression of the HLA-E*01:03 SCT (Figure 4D, left-hand column, top row), but abolished staining with 4D12 (Figure 4D, left-hand column, bottom row). In contrast, 4D12 stained cells expressing the Mamu-E*02:16 SCT with the opposite mutations (Figure 4D, right-hand column, bottom row), despite the three changes apparently causing a reduction in cell surface expression of this SCT (Figure 4D, right-hand column, top row).

When these three positions were mutated individually in the HLA-E*01:03 SCT, only the E144A change had a marked effect on 4D12 staining (Figure 5A, bottom row). 4D12 staining was similarly unaffected by combination of the I142T and S151G mutations (Figure 5B, bottom row, column 2). Despite this, positions 142 and 151 are also important for the binding of 4D12: combination of either I141T or S151G with E144A abolished 4D12 staining (Figure 5B, bottom row, columns 1 and 3, respectively). Interpretation of the effects of the opposite mutations introduced into the Mamu-E*02:16 SCT (Figure 5C and D) was complicated by the fact that some of the mutations again affected cell surface expression of the SCT. However, A144E was the only one of the single mutants to allow 4D12 binding (Figure 5C, bottom row, column 2), and the combination of T142I with G151S was not sufficient to allow 4D12 to bind (Figure 5D, bottom rows, column 2).

**FIGURE 5:**
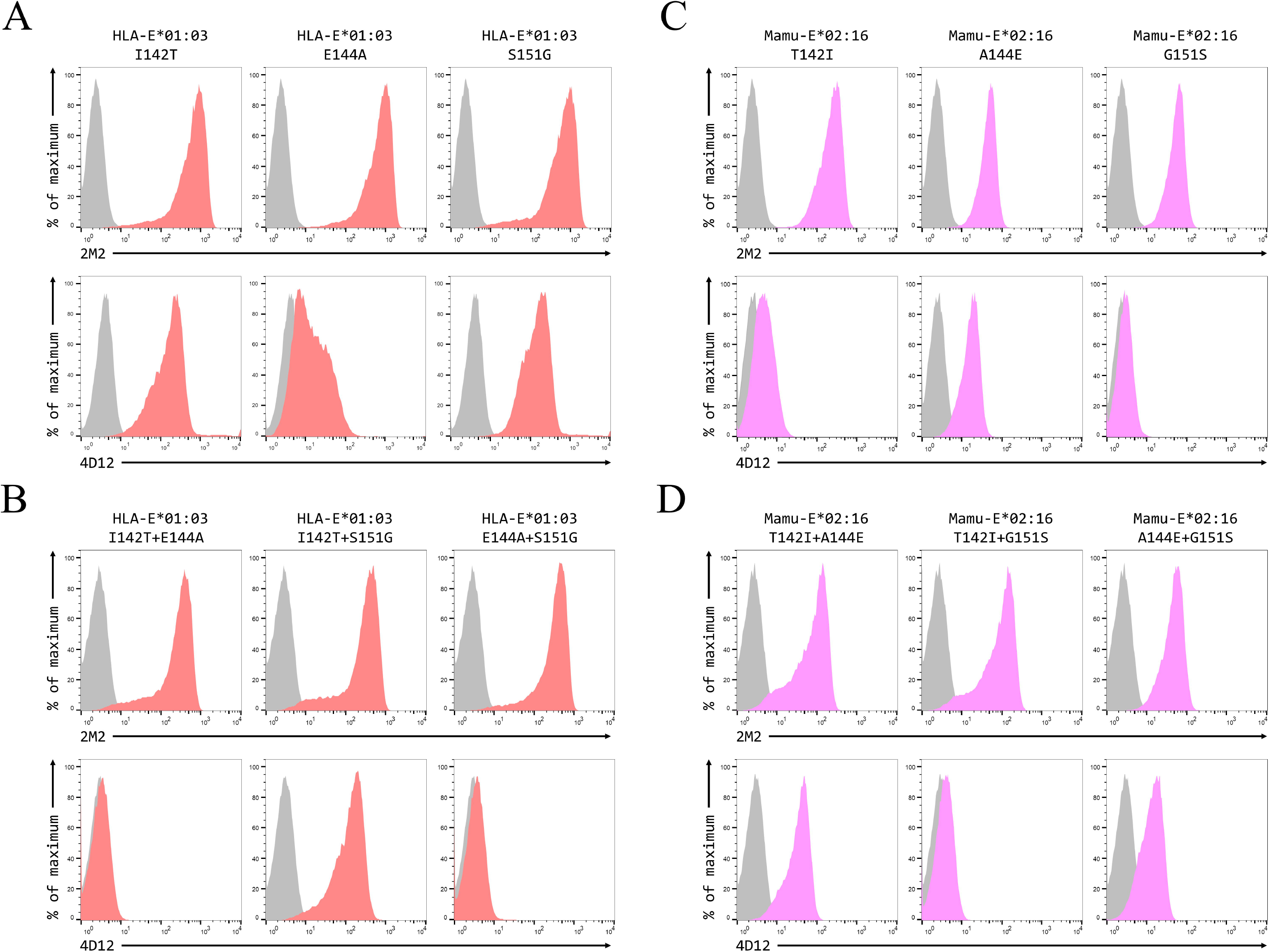
Positions 142, 144 and 151 are all important for the binding of 4D12. Representative staining with 2M2 (top row), or 4D12 (bottom row), of β2-microglobulin-deficient 293T cells transfected with plasmids expressing **(A)** the HLA-E*01:03 SCT with positions 142 (left column), 144 (central column) or 151 (last column) individually mutated to the amino acids present in Mamu-E*02:16; **(B)** the Mamu-E*02:04 SCT with positions 142 (left column), 144 (central column) or 151 (last column) individually mutated to the amino acids present in HLA-E-E*01:03; **(C)** the HLA-E*01:03 SCT with positions 142 and 144 (left column), 142 and 151 (central column) or 144 and 151 (last column) mutated to the amino acids present in Mamu-E*02:16; or **(D)** plasmids expressing the Mamu-E*02:04 SCT with positions 142and 144 (left column), 142 and 151 (central column) or 144 and 151 (last column) mutated to the amino acids present in HLA-E-E*01:03. In all panels, grey histograms show staining of mock transfected cells.

These results confirm the previous observation that 3D12 and 4D12 recognise distinct regions on the HLA-E protein^22^. Moreover, the dramatic effect on 4D12 binding caused by the single and double mutations is consistent with these positions playing a direct role in the binding of 4D12 to what is known to be a conformationally-independent epitope. Although the contributions of the other (conserved) residues in this region remain to be determined, we conclude that the glutamate at position 144 is critical for 4D12 binding, with the isoleucine at position 142 and the serine at position 151 also contributing to the stability of the interaction.

### Cross-reactivity of 4D12 with classical MHC class I alleles

We used a Luminex assay (LABScreen Single Antigen Class I Combi beads, One Lambda) to assess 4D12 binding to 97 individual MHC Ia allotypes (31 HLA-A, 50 HLA-B, and 16 HLA-C; Figure 6A). In addition, we repeated our analysis of the cross-reactivity of 3D12 (Figure 6B). Both antibodies exhibited only weak cross-reaction, although the results with 3D12 differed slightly from what we observed previously^20^ –likely due to batch differences in the beads used.

**FIGURE 6:**
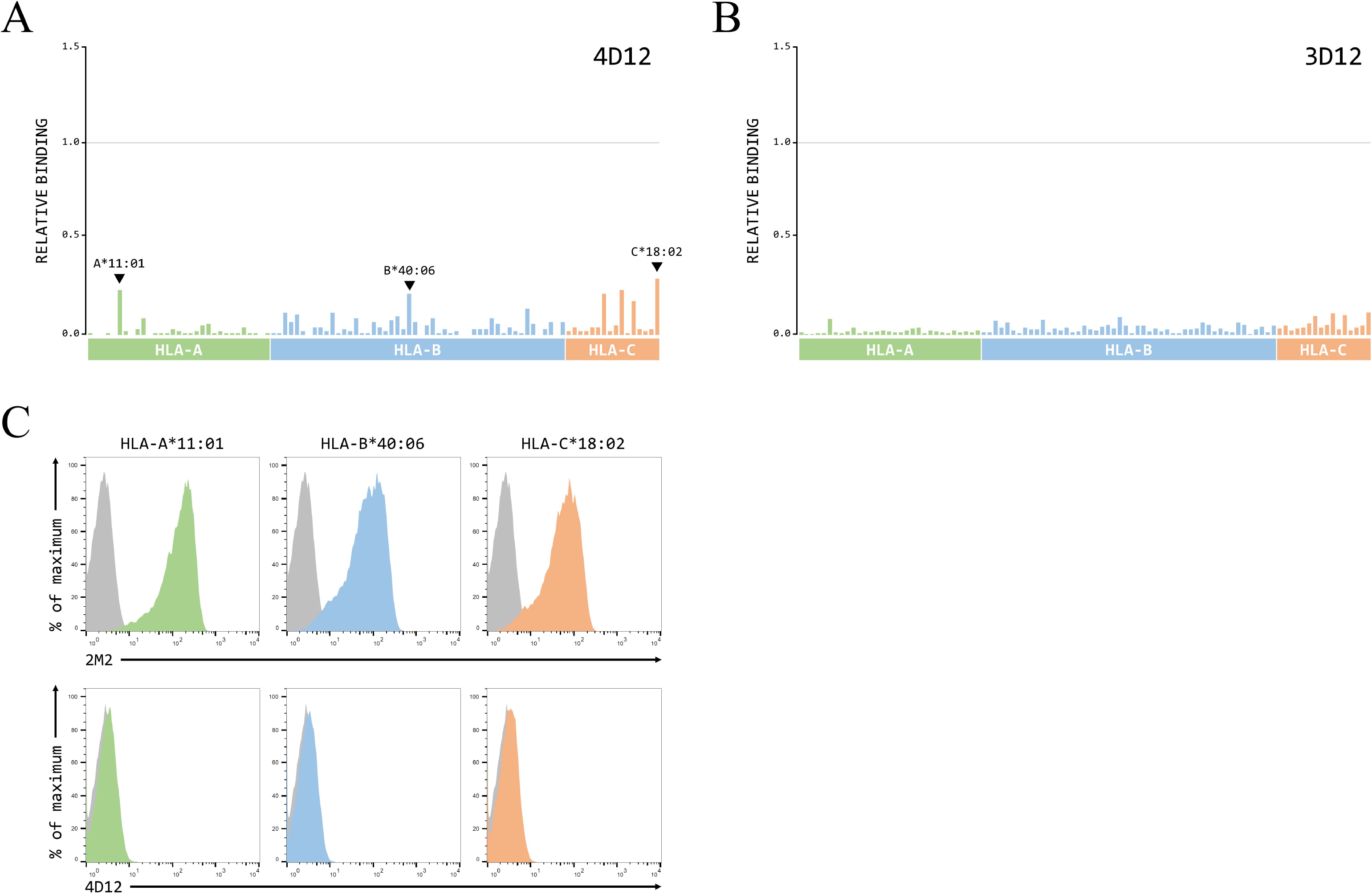
4D12 does not cross-react with classical MHC class I proteins. **(A, B)** Binding of 4D12 (panel A) or 3D12 (panel B) to beads coated with recombinant HLA-A (green), B (blue) and C (red) protein, normalised relative to the binding of the pan-HLA antibody W6/32 (set to 1). The allotypes tested were: HLA-A*01:01, 02:01, 02:03, 02:06, 03:01, 11:01, 11:02, 23:01, 24:02, 24:03, 25:01, 26:01, 29:01, 29:02, 30:01, 30:02, 31:01, 32:01, 33:01, 33:03, 34:01, 34:02, 36:01, 43:01, 66:01, 66:02, 68:01, 68:02, 69:01, 74:01, 80:01; HLA-B*07:02, 08:01, 13:01, 13:02, 14:01 14:02, 15:01, 15:02, 15:03, 15:10, 15:11, 15:12, 15:13, 15:16, 18:01, 27:05, 27:08, 35:01, 37:01, 38:01, 39:01, 40:01, 40:02, 40:06, 41:01, 42:01, 44:02, 44:03, 45:01, 46:01, 47:01, 48:01, 49:01, 50:01, 51:01, 51:02, 52:01, 53:01, 54:01, 55:01, 56:01, 57:01, 57:03, 58:01, 59:01, 67:01, 73:01, 78:01, 81:01, 82:01; HLA-C*01:02, 02:02, 03:02, 03:03, 03:04, 04:01, 05:01, 06:02, 07:02, 08:01, 12:03, 14:02, 15:02, 16:01, 17:01, 18:02. **(C)** Representative staining with 2M2 (top row), or 4D12 (bottom row), of β2-microglobulin-deficient 293T cells transfected with plasmids expressing single chain dimers (SCD) of the three HLA-A, B or C alleles that gave the highest level of cross-reaction on the beads: HLA-A*11:01 (left column), HLA-B*40:06 (central column), and HLA-C*18:01 (last column). Grey histograms show staining of mock transfected cells.

A small number of MHC Ia allotypes gave higher levels of binding with 4D12 than the others, suggesting low lev cross-reactivity. A proportion of the recombinant MHC protein bound to these beads may be mis-folded, potentially leading to higher levels of non-specific binding with particular allotypes. Therefore, to test if the highest levels of cross-reaction seen here would result in 4D12 staining the HLAs expressed on the surface of cells, we transfected β2-miroglobulin-deficient 293T cells with single chain dimers of HLA-A*11:01, HLA-B*40:06 and HLA-C*18:02. All three SCD expressed well at the cell surface (Figure 6C, top row), but none stained with 4D12 (Figure 6C, bottom row). Therefore, we conclude that 4D12 is specific for MHC-E, and does not cross-react with any of the MHC Ia allotypes tested here.

## DISCUSSION

By employing hybrids of a number of different MHC-E alleles, we mapped the epitopes of the most commonly used commercially-available HLA-E antibody, 3D12, as well as the epitope bound by the antibody 4D12, which binds both HLA-E and certain non-human primate MHC-E alleles. Our results confirm the previous observation that these antibodies recognise distinct epitopes; 4D12 binds to the alpha 2 domain of MHC-E, adjacent to the C terminus of the presented peptide, while 3D12 binds on the alpha 3 domain, on the opposite side to that bound by β2-microglobulin, close to the plasma membrane.

The epitope of 3D12 encompasses 4 amino acids that differ between HLA-E and Mamu-E, positions 215, 219, 223 and 224. The position 215 leucine present in HLA-E is conserved in the majority of MHC Ia allotypes, while the amino acids present at the other positions in MHC 1a match those present in all 31 unique Mamu-E proteins in the IPD Non-Human Primates database (release 3.8.0.0) rather than HLA-E (Supplementary Figure 2). Although this might indicate that position 215 does not form part of the epitope, none of the individual mutations of these four positions affected 3D12 binding. The exact contacts made by 3D12 remain to be determined, although precise mapping of these interactions may be hampered by the conformational nature of the 3D12 epitope. The location of the 3D12 epitope is distinct from the sequences identified by a previous attempt to map the epitope of this antibody that employed indirect methods^26^, which identified two sequences (115-QFAYDGKDY-123 and 137-DTAAQI-142), which are conserved not only in Mamu-E*02:04 but also in many HLA-B and C alleles. We note that these authors previously suggested the same two regions for the epitope of the MEM-E/02 antibody ^21^, a conclusion contradicted by a later study that localised the epitope of this antibody to 54-QEGSEYWDRET-64^27^. Such erroneous mapping highlights the difficulty in using indirect assays to map antibody epitopes.

In contrast to the 3D12 epitope, the linear nature of the 4D12 epitope allowed a more detailed mutagenic analysis of the contributions of a number of the alpha 2 domain polymorphisms to the binding of the antibody. The three positions in this region that differ between Mamu-E*02:04 and Mamu-E*02:16 (142, 144, and 151) are all important for binding, with the glutamate at position 144 appearing to be most critical. Further mutagenesis will be required to evaluate the contributions to 4D12 binding of the amino acids that neighbour these polymorphic positions and fully define the exact extent of the epitope.

Although 4D12 has previously been used to stain cells expressing Mamu-E and Mafa-E (rhesus and cynomolgus macaques, respectively) we have shown for the first time that 4D12 does not bind all Mamu-E alleles. In addition to Mamu-E*02:16, six other Mamu-E alleles have changes at positions 142, 144 or 151 (Supplementary Figure 3A), although only two of these have the A144E polymorphism that is most detrimental for 4D12 binding. For Mafa-E (Supplementary Figure 3B), all 14 known alleles (IPD Non-Human Primate database, release 3.8.0.0) have glutamate at 144, and only one allele has a changed key residue (Mafa-E*02:15:01:01, which has glycine at position 151) but would still be expected to be bound by 4D12.

The LABScreen beads assay (Figure 6A) suggested that 4D12 cross-reacted at a low level with certain HLA-A, B and C alleles, but staining of cells expressing SCD of the most cross-reactive alleles was negative (Figure 6C). A comparison of the sequences encompassing the 4D12 epitope in the HLA-A, B and C proteins present on the LABScreen beads (Supplementary Figure 4) showed that, while all alleles have isoleucine at position 142, none have serine at 151 or the critical glutamate at position 144. Moreover, the sequence across the region bound by 4D12 in those alleles that gave the highest 4D12 binding in the LABScreen beads assay is identical to that of alleles that show much less cross-reaction (for example, HLA-A*11:01 and HLA-A*11:02, HLA-B*40:06 and HLA-B*40:02, and HLA-C*18:02 and HLA-C*01:02). It is known that the beads used are coated with a mix of correctly folded and mis-folded protein^33^, so the level of cross-reaction may reflect the amounts of mis-folded protein present on the beads and result from non-specific binding. Such differences in the relative amounts of correctly and incorrectly folded protein most likely explain the slight difference in the 3D12 cross-reactivity profile reported here with that which we published previously^20^.

For 3D12, the fact that the epitope is close to the plasma membrane may mean that its binding may be affected by changes in the lipid composition of the membrane, as observed previously for the pan-MHC class I antibody W6/32^24^. However, 3D12 and 4D12 differ from W6/32 in one important regard: their epitopes comprise only HLA-E sequence, whereas W6/32 recognises a discontinuous epitope involving residues in both the HLA and β2-microglobulin^34^. For MHC Ia, staining with W6/32 is likely to be limited to correctly folded complexes, as these are known to rapidly dissociate upon loss of the presented peptide^35^. In contrast, we have previously shown that HLA-E appears to be unusually stable in the absence of peptide in vitro, and that these HLA-E-β2-microglobulin dimers may adopt a range of different conformations^28^. If peptide-free HLA-E has similar stability at the cell surface, this would have important implications for the use of either 3D12 or 4D12 to assess the expression of HLA-E/β2-microglobulin/peptide complexes by cells. As 4D12 recognises a linear epitope, it would not be expected to discriminate between correctly folded HLA-E/β2-microglobulin/peptide complexes and mis-folded or peptide-free HLA-E complexes. For 3D12, the conformational nature of the epitope may impose an element of selectivity in terms of which the forms of HLA-E to which the antibody would be capable of binding. It should be stressed, however, that 3D12 may also be able to bind mis-folded, or even β2-microglobulin-free, complexes, depending on the extent to which its epitope is distorted. Consistent with this, it has been reported that 3D12 can be used for western blotting^36^, although it is possible that this could be explained by incomplete denaturation, or transient renaturation, of the 3D12 epitope depending on the blotting conditions used. Further work on the exact range of complexes that 3D12 and 4D12 can recognise will be essential to fully determine the utility of these antibodies for monitoring cell surface expression of correctly folded HLA-E/β2-microglobulin/peptide complexes.

Despite these concerns about the exact form(s) of HLA-E recognised by 3D12, we have previously used 3D12 as the capture antibody in a sandwich ELISA to measure peptide binding by HLA-E^37^. In this assay, the amount of HLA-E/β2-microgobulin/peptide complex generated in a micro-scale refold, captured by 3D12, and detected using an antibody against β2-microglobulin, depends on the affinity of HLA-E for the peptide. Attempts to set up the same ELISA for Mamu-E using 4D12 have been unsuccessful (G. Gillespie, L. Walters and A.J.M., unpublished data), perhaps because the location of the 4D12 epitope on the alpha 2 domain results in the Mamu-E/β2-microglobulin/peptide complex being held on the ELISA plate in an orientation that hampers access of the anti-β2-microglobulin detection antibody. Even if 4D12 worked as a capture antibody, however, the fact that it recognises a linear epitope would likely compromise the ELISA results, as the captured Mamu-E complexes would be a mix of correctly and incorrectly folded complexes. The same potential problem also exists with a recently published Mamu-E peptide binding ELISA^38^, which used streptavidin to capture biotinylated Mamu-E. To overcome the lack of a suitable Mamu-E-specific capture antibody, we plan to trial a peptide binding ELISA using our M23/H4/M4 hybrid. Although this contains 4 amino acids changes compared with Mamu-E*02:04, these mutations are all on the alpha 3 domain, and would be expected to have little or no impact on peptide binding. If successful, it will be informative to extend it to other alleles of Mamu-E to determine if the polymorphic nature of this MHC has any impact on its peptide repertoire.

In conclusion, we have determined the locations on HLA-E of the epitopes bound by the antibodies 3D12 and 4D12, although further experiments are necessary to elucidate the exact forms of HLA-E that are recognised by both of these antibodies.

## ACKNOWLEDGEMENTS

We thank Andrew Worth for cell sorting, Tim Rostron and John Frankland of the Weatherall Institute of Molecular Medicine for plasmid sequencing, and Geraldine Gillespie as well as past and present members of the McMichael, Gillespie and Borrow laboratories for helpful discussions while this work was being undertaken. The work was supported by funding from the Bill and Melinda Gates Foundation (OPP1133649), the NIH, NIAID, Division of AIDS (Collaboratory of AIDS Researchers for Eradication [CARE]; UM1 AI 126619 and UM1 AI 164567), and the Chinese Academy of Medical Sciences (CAMS) Innovation Fund for Medical Sciences (CIFMS 2018-I2M-2-002). P.B. and A.J.M. are Jenner Institute Investigators.

## AUTHOR CONTRIBUTIONS

S.B. conceived and performed the experiments and analysed the data. P.B. and A.J.M. supervised the work and analysed data. S.B. wrote the manuscript, which was reviewed, edited and approved by all authors.

## REFERENCES

1. Rodgers JR, Cook RG. MHC class Ib molecules bridge innate and acquired immunity. Nat Rev Immunol. Jun 2005;5(6):459–71. doi:10.1038/nri1635

2. Olieslagers TI, Voorter CEM, Groeneweg M, Xu Y, Wieten L, Tilanus MGJ. New insights in HLA-E polymorphism by refined analysis of the full-length gene. Hla. Mar 2017;89(3):143–149. doi:10.1111/tan.12965

3. Strong RK, Holmes MA, Li P, Braun L, Lee N, Geraghty DE. HLA-E allelic variants. Correlating differential expression, peptide affinities, crystal structures, and thermal stabilities. J Biol Chem. Feb 14 2003;278(7):5082–90. doi:10.1074/jbc.M208268200

4. Braud V, Jones EY, McMichael A. The human major histocompatibility complex class Ib molecule HLA-E binds signal sequence-derived peptides with primary anchor residues at positions 2 and 9. Eur J Immunol. May 1997;27(5):1164–9. doi:10.1002/eji.1830270517

5. Braud VM, Allan DS, Wilson D, McMichael AJ. TAP- and tapasin-dependent HLA-E surface expression correlates with the binding of an MHC class I leader peptide. Curr Biol. Jan 1 1998;8(1):1–10. doi:10.1016/s0960-9822(98)70014-4

6. Lee N, Goodlett DR, Ishitani A, Marquardt H, Geraghty DE. HLA-E surface expression depends on binding of TAP-dependent peptides derived from certain HLA class I signal sequences. J Immunol. May 15 1998;160(10):4951–60.

7. Braud VM, Allan DS, O’Callaghan CA, et al. HLA-E binds to natural killer cell receptors CD94/NKG2A, B and C. Nature. Feb 19 1998;391(6669):795–9. doi:10.1038/35869

8. Borrego F, Ulbrecht M, Weiss EH, Coligan JE, Brooks AG. Recognition of human histocompatibility leukocyte antigen (HLA)-E complexed with HLA class I signal sequence-derived peptides by CD94/NKG2 confers protection from natural killer cell-mediated lysis. J Exp Med. Mar 2 1998;187(5):813–8. doi:10.1084/jem.187.5.813

9. Lee N, Llano M, Carretero M, et al. HLA-E is a major ligand for the natural killer inhibitory receptor CD94/NKG2A. Proc Natl Acad Sci U S A. Apr 28 1998;95(9):5199–204. doi:10.1073/pnas.95.9.5199

10. Grant EJ, Nguyen AT, Lobos CA, Szeto C, Chatzileontiadou DSM, Gras S. The unconventional role of HLA-E: The road less traveled. Mol Immunol. Apr 2020;120:101–112. doi:10.1016/j.molimm.2020.02.011

11. Hansen SG, Piatak M, Jr., Ventura AB, et al. Immune clearance of highly pathogenic SIV infection. Nature. Oct 3 2013;502(7469):100-4. doi:10.1038/nature12519

12. Hansen SG, Sacha JB, Hughes CM, et al. Cytomegalovirus vectors violate CD8+ T cell epitope recognition paradigms. Science. May 24 2013;340(6135):1237874. doi:10.1126/science.1237874

13. Hansen SG, Wu HL, Burwitz BJ, et al. Broadly targeted CD8(+) T cell responses restricted by major histocompatibility complex E. Science. Feb 12 2016;351(6274):714-20. doi:10.1126/science.aac9475

14. Malouli D, Hansen SG, Hancock MH, et al. Cytomegaloviral determinants of CD8(+) T cell programming and RhCMV/SIV vaccine efficacy. Sci Immunol. Mar 25 2021;6(57)doi:10.1126/sciimmunol.abg5413

15. Hansen SG, Zak DE, Xu G, et al. Prevention of tuberculosis in rhesus macaques by a cytomegalovirus-based vaccine. Nat Med. Feb 2018;24(2):130–143. doi:10.1038/nm.4473

16. Burwitz BJ, Hashiguchi PK, Mansouri M, et al. MHC-E-Restricted CD8(+) T Cells Target Hepatitis B Virus-Infected Human Hepatocytes. J Immunol. Apr 15 2020;204(8):2169–2176. doi:10.4049/jimmunol.1900795

17. Ogg G, Cerundolo V, McMichael AJ. Capturing the antigen landscape: HLA-E, CD1 and MR1. Curr Opin Immunol. Aug 2019;59:121-129. doi:10.1016/j.coi.2019.07.006

18. Ottenhoff THM, Joosten SA. Mobilizing unconventional T cells. Science. Oct 18 2019;366(6463):302-303. doi:10.1126/science.aay7079

19. Voogd L, Ruibal P, Ottenhoff THM, Joosten SA. Antigen presentation by MHC-E: a putative target for vaccination? Trends Immunol. Mar 31 2022;doi:10.1016/j.it.2022.03.002

20. Corrah TW, Goonetilleke N, Kopycinski J, et al. Reappraisal of the relationship between the HIV-1-protective single-nucleotide polymorphism 35 kilobases upstream of the HLA-C gene and surface HLA-C expression. J Virol. Apr 2011;85(7):3367–74. doi:10.1128/JVI.02276-10

21. Ravindranath MH, Taniguchi M, Chen CW, et al. HLA-E monoclonal antibodies recognize shared peptide sequences on classical HLA class Ia: relevance to human natural HLA antibodies. Mol Immunol. Feb 2010;47(5):1121–31. doi:10.1016/j.molimm.2009.10.024

22. Ishitani A, Sageshima N, Lee N, et al. Protein expression and peptide binding suggest unique and interacting functional roles for HLA-E, F, and G in maternal-placental immune recognition. J Immunol. Aug 1 2003;171(3):1376–84. doi:10.4049/jimmunol.171.3.1376

23. Wu HL, Wiseman RW, Hughes CM, et al. The Role of MHC-E in T Cell Immunity Is Conserved among Humans, Rhesus Macaques, and Cynomolgus Macaques. J Immunol. Jan 1 2018;200(1):49–60. doi:10.4049/jimmunol.1700841

24. Jongsma MLM, de Waard AA, Raaben M, et al. The SPPL3-Defined Glycosphingolipid Repertoire Orchestrates HLA Class I-Mediated Immune Responses. Immunity. Jan 12 2021;54(1):132–150 e9. doi:10.1016/j.immuni.2020.11.003

25. Zhang T, de Waard AA, Wuhrer M, Spaapen RM. The Role of Glycosphingolipids in Immune Cell Functions. Front Immunol. 2019;10:90. doi:10.3389/fimmu.2019.00090

26. Ravindranath MH, Pham T, El-Awar N, Kaneku H, Terasaki PI. Anti-HLA-E mAb 3D12 mimics MEM-E/02 in binding to HLA-B and HLA-C alleles: Web-tools validate the immunogenic epitopes of HLA-E recognized by the antibodies. Mol Immunol. Jan 2011;48(4):423–30. doi:10.1016/j.molimm.2010.09.011

27. Tremante E, Lo Monaco E, Ingegnere T, Sampaoli C, Fraioli R, Giacomini P. Monoclonal antibodies to HLA-E bind epitopes carried by unfolded beta2 m-free heavy chains. Eur J Immunol. Aug 2015;45(8):2356–64. doi:10.1002/eji.201545446

28. Walters LC, Harlos K, Brackenridge S, et al. Pathogen-derived HLA-E bound epitopes reveal broad primary anchor pocket tolerability and conformationally malleable peptide binding. Nat Commun. Aug 7 2018;9(1):3137. doi:10.1038/s41467-018-05459-z

29. Higuchi R, Krummel B, Saiki RK. A general method of in vitro preparation and specific mutagenesis of DNA fragments: study of protein and DNA interactions. Nucleic Acids Res. Aug 11 1988;16(15):7351–67. doi:10.1093/nar/16.15.7351

30. Ren J, Liu X, Fang C, Jiang S, June CH, Zhao Y. Multiplex Genome Editing to Generate Universal CAR T Cells Resistant to PD1 Inhibition. Clin Cancer Res. May 1 2017;23(9):2255–2266. doi:10.1158/1078-0432.CCR-16-1300

31. Perez-Pinera P, Kocak DD, Vockley CM, et al. RNA-guided gene activation by CRISPR-Cas9-based transcription factors. Nat Methods. Oct 2013;10(10):973–6. doi:10.1038/nmeth.2600

32. Ding Q, Regan SN, Xia Y, Oostrom LA, Cowan CA, Musunuru K. Enhanced efficiency of human pluripotent stem cell genome editing through replacing TALENs with CRISPRs. Cell Stem Cell. Apr 4 2013;12(4):393–4. doi:10.1016/j.stem.2013.03.006

33. Jucaud V, Ravindranath MH, Terasaki PI. Conformational Variants of the Individual HLA-I Antigens on Luminex Single Antigen Beads Used in Monitoring HLA Antibodies: Problems and Solutions. Transplantation. Apr 2017;101(4):764–777. doi:10.1097/TP.0000000000001420

34. Ladasky JJ, Shum BP, Canavez F, Seuanez HN, Parham P. Residue 3 of beta2-microglobulin affects binding of class I MHC molecules by the W6/32 antibody. Immunogenetics. Apr 1999;49(4):312–20. doi:10.1007/s002510050498

35. Ljunggren HG, Stam NJ, Ohlen C, et al. Empty MHC class I molecules come out in the cold. Nature. Aug 2 1990;346(6283):476-80. doi:10.1038/346476a0

36. Lo Monaco E, Sibilio L, Melucci E, et al. HLA-E: strong association with beta2-microglobulin and surface expression in the absence of HLA class I signal sequence-derived peptides. J Immunol. Oct 15 2008;181(8):5442–50. doi:10.4049/jimmunol.181.8.5442

37. Walters LC, McMichael AJ, Gillespie GM. Detailed and atypical HLA-E peptide binding motifs revealed by a novel peptide exchange binding assay. Eur J Immunol. Dec 2020;50(12):2075–2091. doi:10.1002/eji.202048719

38. Ruibal P, Franken K, van Meijgaarden KE, et al. Peptide Binding to HLA-E Molecules in Humans, Nonhuman Primates, and Mice Reveals Unique Binding Peptides but Remarkably Conserved Anchor Residues. J Immunol. Nov 15 2020;205(10):2861–2872. doi:10.4049/jimmunol.2000810

